# The Rate of Molecular Evolution When Mutation May Not Be Weak

**DOI:** 10.1101/259507

**Authors:** A.P. Jason de Koning, Bianca D. De Sanctis

## Abstract

One of the most fundamental rules of molecular evolution is that the rate of neutral evolution equals the mutation rate and is independent of effective population size. This result lies at the heart of the Neutral Theory, and is the basis for numerous analytic approaches that are widely applied to infer the action of natural selection across the genome and through time, and for dating divergence events using the molecular clock. However, this result was derived under the assumption that evolution is strongly mutation-limited, and it has not been known whether it generalizes across the range of mutation pressures or the spectrum of mutation types observed in natural populations. Validated by both simulations and exact computational analyses, we present a direct and transparent theoretical analysis of the Wright-Fisher model of population genetics, which shows that some of the most important rules of molecular evolution are fundamentally changed by considering recurrent mutation’s full effect. Surprisingly, the rate of the neutral molecular clock is found to have population-size dependence and to not equal the mutation rate in general. This is because, for increasing values of the population mutation rate parameter (*θ*), the time spent waiting for mutations quickly becomes smaller than the cumulative time mutants spend segregating before a substitution, resulting in a net deceleration compared to classical theory that depends on the population mutation rate. Furthermore, selection exacerbates this effect such that more adaptive alleles experience a greater deceleration than less adaptive alleles, introducing systematic bias in a wide variety of methods for inferring the strength and direction of natural selection from across-species sequence comparisons. Critically, the classical weak mutation approximation performs well only when *θ<* 0.1, a threshold that many biological populations seem to exceed.

## Introduction

Classical population genetic theory was largely based on the assumption that mutation is “weak” and can therefore be conveniently ignored. Although it has been fairly clear that this assumption is often reasonable, estimates of the population mutation rate parameter (*θ*; see Methods) across many organisms have only become available recently in the genomics era, and it has been unclear what values of *θ* might violate weak mutation. A variety of examples are now known of populations, and mutation types, where mutation might very well not be weak. These include hyperdiverse eukaryotes (Cutter *et al*., 2013), many prokaryotes (Hughes *et al*., 2008; Sung *et al*., 2012), and a variety of rapidly evolving viruses (including HIV; Maldarelli *et al*., 2013; Pennings, Kryazhimskiy, and Wakeley, 2014; Rouzine, Coffin, and Weinberger, 2014). Similarly, mutation types with fast natural rates such as some context-dependent nucleotide mutations in the nuclear genomes of mammals (Ehrlich and Wang, 1981), mutations in the mitochondrial genomes of some vertebrates (Brown *et al*., 1979), microsatellite and simple sequence repeat polymorphisms (Richard *et al*., 2008), somatic mutations (Lynch, 2010), and heritable epigenetic changes (Charlesworth and Jain, 2014) all occur at rates fast enough that it is reasonable to question the correctness of the weak mutation assumptions upon which most analytic approaches rely. Furthermore, as has been recently pointed out (Messer *et al*., 2016), the apparent ubiquity of “soft sweeps” in nature (Karasov *et al*., 2010; Messer and Petrov, 2013; Pennings and Hermisson, 2005, 2006), where adaptive mutations appear to have multiple origins by recurrent mutation or immigration, has been interpreted as supporting the idea that *θ* in some populations may be significantly larger than is widely believed. It is therefore critical to determine how the implications of classical population genetic theory might change under the degrees of mutation pressure observed in natural populations (Charlesworth and Jain, 2014; Karasov *et al*., 2010; Kryazhimskiy and Plotkin, 2008; Messer *et al*., 2016; Pennings and Hermisson, 2006).

Here we explore the impact of general, recurrent mutation processes on the rate of molecular evolution. The rate of evolution is among the most fundamental and useful quantities in all of evolutionary genetics, and is the basis for analytic approaches used widely in the study of molecular evolution, genome evolution, population genetics, phylogenetics, and related fields. Despite its central importance to understanding patterns of genomic sequence variations and their causes, remarkably, no explicit and complete derivation has appeared in the literature. Kimura (1968), like Wright (1938), seems to have simply written down the standard equation from intuition, and it has since become second nature to population geneticists. However, this rate of evolution depends upon several important implicit assumptions that appear to not be widely appreciated. Here we make those assumptions explicit, and after doing so, show that it is surprisingly easy and valuable to generalize the rate of evolution at individual genomic positions with respect to mutation.

Sequence evolution is often considered from two rather different perspectives. When making across-species sequence comparisons, molecular evolution is typically modelled at individual genomic positions following a phylogenetic *substitution process*, wherein recurrent mutations occur over long timescales and evolution proceeds by successive substitution events, at the same positions, along diverging lineages. At the population genetic level, each substitution corresponds to the turnover of the entire population for a new allelic state. It has long been considered essential to account for the possibility of serial substitutions at the same positions (e.g., Felsenstein 2003; Yang 2014). Indeed, standard phylogenetic likelihood computations allow for an infinity of unobservable serial substitutions, sometimes represented as a distribution of substitution histories (reviewed in de Koning, Gu, and Pollock, 2009). This is the canonical approach that is implicitly applied when using continuous-time Markov chain models of sequence evolution.

Working from a rather different perspective, Kimura was among the first to consider how the rate of molecular evolution could be approximated in terms of the underlying population genetic processes that generate substitutions (e.g., Kimura 1968; King and Jukes 1969; Wright 1938; also see Bustamante 2005 for an excellent review). In particular, he considered the rate of long-term evolution by fixation under free recombination, no epistasis, and weak mutation. We will refer to this quantity as the *weak-mutation rate of evolution* (defined and discussed in detail in the next section). Kimura justified his approach by referring to the assumptions of the infinite-sites model, which assumes weak mutation that can never be recurrent, although he does not appear to have claimed that his approach requires it (Kimura, 1983, p.46). Although the infinite-sites assumption should generally preclude the application of Kimura’s model to substitution processes at individual positions, in practice his model and its insights are widely applied to individual sites. For example, they are applied implicitly in *d*_*N*_ */d*_*S*_ approaches for inferring the strength and direction of natural selection (Goldman and Yang, 1994; Messier and Stewart, 1997; Muse and Gaut, 1994; Yang, 1998; Yang *et al*., 2000; Zhang *et al*., 2005), and explicitly in methods for inferring population-genetic parameters from across-species comparisons, such as in so called mutation-selection models of codon substitution (Dimmic *et al*., 2000; Halpern and Bruno, 1998; Nielsen and Yang, 2003; Rodrigue *et al*., 2010; Thorne *et al*., 2007). As we discuss in detail below, this apparent contradiction is explained by noting that Kimura’s derivation actually does not require infinite-sites *per se*, but rather it requires some specific, related assumptions about the weakness of mutation.

Before we derive the rate of evolution under general mutation, some potentially counter-intuitive ideas must first be introduced. Owing to variation in definitions of a fixation, substitution and fixation may or may not correspond. Fixation is most often defined as the takeover of the entire population by a single mutant lineage. This is a convenient definition because it has an approximate inverse correspondence to coalescence. However, in standard models of population genetics, such as the Wright-Fisher model, fixation is usually defined as simply reaching 100% frequency for the mutant *state*. These definitions coincide solely when mutation is unnaturally disallowed in the instantaneous generator of the underlying model, so that only a single lineage of mutants is permitted to exist in the population at a time. Although there are reasons to be interested in fixations that leave the population identical-by-descent (IBD), this particular approach is undesirable because it simply labels all mutants as IBD by assumption, ignoring the possibility of additional mutations.

Even over short timescales, this can be highly unrealistic. For example, neutral mutations that go to fixation persist on average for 4*N*_*e*_ generations in diploid populations, where *N*_*e*_ is the effective population size and may differ from the census population size, *N*. Since approximately 2*Nv* mutations are expected to arise in each generation, for forward mutation rate *v*, the number of additional mutations expected in the population during an average neutral fixation trajectory exceeds 1 when the mutation rate is as small as *v* = 1*/*(8*NN*_*e*_), or 1*/*(8*N* ^2^) when *N*_*e*_ = *N*. Thus, even for exceedingly small mutation rates, it is plausible that multiple lineages of the same variant commonly arise simultaneously. When population mutation rates are relatively large, it is substantially more likely that this will occur (e.g., soft selective sweeps; Pennings and Hermisson, 2005). This is important because the standard, weak-mutation rate of evolution considers only the fixation of lineages with single mutational origins. However, especially when making across-species sequence comparisons where we typically consider only single sequences from each species, we generally assume that so many generations have elapsed that positions evolved independently via effectively free recombination. We therefore cannot necessarily tell whether an apparent substitution had single or multiple mutational origins when measuring the rate of long-term evolution. Rather, all we can confidently infer from phylogenetic comparisons is that a position has turned over for a new state. Therefore, we argue that the rate of molecular evolution in across-species sequence comparisons should correspond to the rate of substitution by either single *or* multiple mutational origins. When mutation is weak, this substitution rate will be appropriately dominated by fixations with single origins, whereas when it is stronger, fixations with multiple origins will matter more.

### The rate of evolution under weak mutation

Kimura (1968, 1983), and King and Jukes (1969), building on earlier work by Wright (1938), first showed that the rate of neutral evolution is expected to equal the mutation rate when back mutation is ignored. To obtain this result, they started with an expression for the diploid *weak-mutation rate of evolution*,

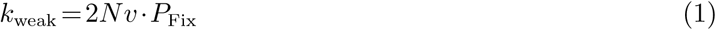

where *N* is the number of reproducing individuals, *v* the forward mutation rate per locus per generation, and *P*_Fix_ the probability that a mutation will eventually go to fixation (see McCandlish and Stoltzfus, 2014, for a perspective on the history of such ‘origin-fixation’ models.) Consistent with common practices described above, we define a locus as an individual genomic position or site. Since the probability of fixation for neutral mutations is 1/(2*N*) under weak mutation assumptions (discussed below), the *weak-mutation rate of neutral evolution* is

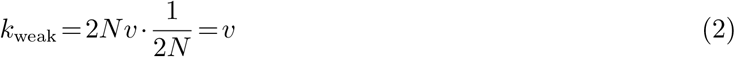

This result is a cornerstone principle of the Neutral Theory of Molecular Evolution (Kimura, 1983) and is deeply embedded in our thinking about the relationship between population genetics and molecular evolution (but see Allen *et al*., 2015; Lanfear *et al*., 2014, for possible exceptions in non-equilibrium populations.) Indeed, it has been called “one of the most elegant and widely applied results in population genetics” (Draghi *et al*., 2011). Although Kimura described this equation in the context of the infinite-sites model (Kimura, 1983, p. 46), it is just as consistent when interpreted in a finite-sites context where mutation is assumed to be weak. Indeed, as we argued above, it is this context in which equation 1 is usually applied. However, as we will show, relaxing the weak mutation assumptions used to derive this result leads to a different rate of evolution, which can have strikingly different characteristics.

To appreciate why equation 1 must require some assumptions about weak-mutation, it is useful to consider the implications of equation 2 if applied without caveats. Naive application of equation 2 predicts that the rate of neutral evolution increases linearly and *without bound* as the mutation rate is increased. However, variants do not reach their fates instantaneously. Therefore, when mutation is fast enough that the time it takes to get a mutation in the population is of the same order as the time it takes for variants to reach their fates, segregation times are expected to at least partly determine the rate of evolution. This suggests that the rate of evolution might level off for increasing mutation rates. To understand what caveats should thus be applied, it is useful to consider the assumptions built into equation 1.

### The weak mutation assumptions

Several assumptions about weak mutation are implied by equation 1. These are: 1) that mutations arise and go to their fates one by one, so that only one segregating lineage of mutations may exist in a population at a given time; 2) that evolution is fundamentally mutation-limited, so that the timescale of mutation dominates over segregation times (Crow and Kimura, 1970); and 3) that mutations originate in a single individual at a time (i.e., the initial number of mutant alleles is *p* = 1).

When mutation is very weak, these assumptions are reasonable because evolution is strongly mutation-limited and populations will be in what has been called the “successional-mutations regime” (Desai and Fisher, 2007). When mutation is not weak, however, these assumptions lead to strange and unrealistic consequences if retained (e.g., in the “concurrent-mutations regime”; Desai and Fisher, 2007). For example, by assuming that mutations arise and reach their fates one-at-a-time (weak-mutation assumption 1), mutations are allowed only when the population is monomorphic. Thus, mutation is unrealistically allowed and then disallowed according to the frequency of the mutant state in the population. As a result, soft sweeps can’t happen, since only one lineage of the mutant state is allowed in the population by assumption. Also when mutation is not weak, the waiting time for mutations per fixation may no longer exceed the time it takes for mutants to reach their fates, thus violating weak mutation assumption 2. In discussing this assumption and its relation to the rate of neutral evolution, Crow and Kimura (1970, p. 369) claimed that the waiting time for mutations should be safely larger than the *fixation time* as long as *θ ≪* 1. This criterion is so broad, however, as to provide almost no intuition about how small *θ* must be for the weak-mutation assumptions to be met. Furthermore, as we will show, it is not just the fixation time that needs to be considered.

Recent work on the effects of arbitrarily fast mutation (Charlesworth and Jain, 2014) has retained the weak-mutation assumptions, perhaps due to the perception that they are needed for analytic tractability. Contrariwise, we will first show how the first two assumptions can be easily relaxed without approximation, and will turn our attention to the third in the supplementary methods (SI Methods 1.1). Throughout this work, we assume a biallelic locus undergoing recurrent bidirectional mutation. Because we are interested in when the mutant *state* takes over the population (e.g., by either a hard or a soft sweep), we do not distinguish between individuals who are identical by state or identical by descent. Importantly, this means that the usual inverse correspondence between fixation and coalescence is lost, except when *θ* is very small (see Methods for full details).

## Results

### The rate of evolution

Following Kimura (1983), we define the rate of evolution as one over the mean time between substitutions. The rate is thus measured in expected substitutions per generation. This time can be equivalently expressed as the mean time to the first substitution, starting at *t* = 0 with a population that is 100% wildtype. For simplicity, we will refer to this interval as the ‘substitution cycle’. In the finite-sites context, this rate of evolution refers to the rate of substitution of different allelic states at the same position. Guess and Ewens (1972) referred to this as the rate of “quasifixation”, however, we will avoid this term since others have used it to mean something different. Despite our rejection of this term, it is worth recognizing the appropriateness of Guess and Ewens’ intent, which seeks to clarify that ‘fixation’ imperfectly describes what happens in real populations where, with recurrent mutation, nothing is every really ‘fixed’ (*i*.*e*., except by purifying selection).

Except where noted otherwise, we will henceforth use the term substitution to refer to the replacement of a population monomorphic for the wildtype state with a population monomorphic for the the mutant state (involving either single or multiple origins). Because fixations are rare, even for advantageous mutations (Kimura and Ota, 1971), for every mutation that arises and becomes fixed, we expect many more mutations to have arisen and gone extinct. We call these mutation-fixation (MF) and mutation-extinction (ME) cycles respectively (or mutation-absorption cycles, in the general case). One over the mean length of the substitution cycle can then be written as a function of the expected number of mutation-absorption cycles and their respective lengths,

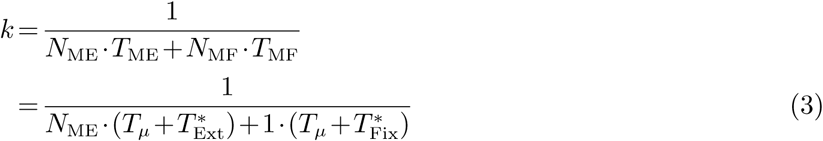

where *N*_*M·*_ and *T*_*M·*_ denote the mean number of cycles and the mean length of cycles, respectively. *T*_*µ*_, 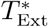, and 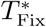 are defined as the mean time in numbers of generations to get a mutation (or mutations), the mean time to extinction (calculated including the effect of bidirectional mutation, as indicated by the ‘*’), and the mean time to fixation (also including mutation).

Since 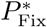 represents the probability that an absorption is a fixation, 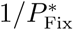 is the expected number of absorptions to get a fixation. Note that each absorption first requires a mutation, and each trial thus represents an originating mutation and its absorption. Since one of these will be a mutation-fixation cycle, 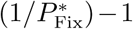 of these are expected to be mutation-extinction cycles. We can therefore write

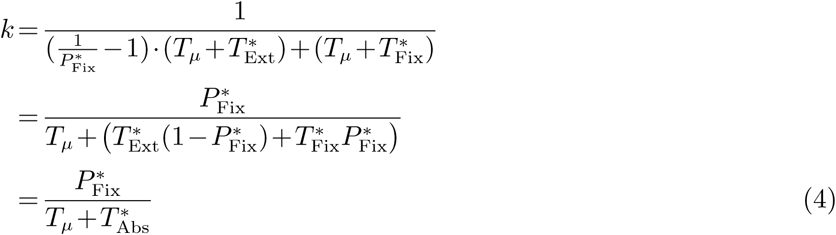

where 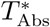 is the unconditional time to absorption allowing for bidirectional mutation. We will refer to equation 4 simply as the rate of evolution, in contrast to the “weak-mutation rate of evolution” (equation 1). Notably, by reintroducing Kimura’s assumptions into equation 4, this expression becomes equal to equation 1 (SI Methods 1.2). We also provide a more formal derivation of equation 4 in the supplement (SI Methods 1.3), and show how it can be integrated over an initial distribution, *f* (*p*) (SI Methods 1.1), thus relaxing the third weak mutation assumption that *p* = 1. All subsequent results are integrated over *f* (*p*).

Unlike the weak-mutation rate of evolution, equation 4 makes no assumptions about the strength of mutation and indeed introduces no additional assumptions beyond those of the model of population genetics used to calculate its component quantities. Dominance, selection, recurrent bidirectional mutation, and other forces may thus be easily considered by including their effects in the underlying model. Importantly, simply incorporating mutation into the probability of fixation in equation 1 does not work without also including the absorption times (Fig. S1). Although simple closed-form expressions for the component quantities are not available using diffusion theory, they can be easily calculated directly from the underlying Markov model using efficient computational techniques we recently described (De Sanctis *et al*., 2017; Krukov *et al*., 2016).

### Direct computation of the rate of evolution

For validation, we developed a direct way to compute the mean time between fixations, without requiring any of the above theory. This direct approach uses a modified Wright-Fisher model, where the extinction state is treated as transient rather than absorbing, and the population is initialized with *p* = 0 mutants. This allows the time between fixations to be directly calculated as the expected time to absorption, using standard absorbing Markov chain theory (Krukov *et al*., 2016, see Methods). A similar approach can be used to directly calculate the variance of the time between fixations, which is useful for testing hypotheses about the dispersion of the molecular clock (Fig. S3). When equation 4 is integrated over *f* (*p*), it numerically agrees with the direct approach (see Methods; Fig. S2).

### The weak-mutation assumptions are violated for surprisingly small population mutation rates

To first clarify the range of population mutation rates where the weak-mutation assumptions are valid, we conducted neutral Wright-Fisher simulations under both one-at-a-time mutation scenarios (Simulation I), and under a uniform mutation model where mutations can arise naturally in any generation according to the mutation rate (Simulation II). For these and most of the other analyses to follow, we set the back mutation rate to zero to be conservative in our assessment of how the full rate of evolution differs from the weak-mutation rate of evolution (since the latter does not include back-mutation; also see Methods). This means that the results to follow are most directly applicable to types of mutations that are unidirectionally fast (e.g., transitions at methylated CpG’s in mammals). However, the rate of evolution with bidirectional mutation is addressed at the end of the Results section, where our main findings are seen to hold and, expectedly, to be even more extreme.

To first demonstrate and verify that the one-at-a-time mutation model is consistent with the assumptions of equation 2, as we have claimed, we compared simulation averages for several quantities to their expectations under the weak-mutation rate of neutral evolution. Based on equation 2, the expected number of trials (i.e., mutation-absorptions) to get a fixation is 1*/P*_Fix_ = 2*N*, and the expected amount of time spent waiting for mutations per fixation, is 1*/P*_Fix_ *·T*_*µ*_ = 1*/v* (assuming Poisson-distributed mutations). By the second weak-mutation assumption, the expected time between fixations is then simply the expected time spent waiting for mutations per fixation (or 1*/v* generations).

Simulation averages for the number of trials to get a fixation were tightly clustered around 2*N* regardless of the mutation rate (Table 1, Simulation I), and the mean time spent waiting for mutations per fixation was similarly tightly clustered around the expected value of 1*/v* generations. Therefore, if we assume the mean time spent waiting for mutations per fixation closely approximates the expected time between fixations, the simulation results agree precisely with the predictions of equation 1. Notice, however, that in these simulations we can also directly examine the total amount of time between fixations without invoking the second weak-mutation assumption.

**Table 1.**
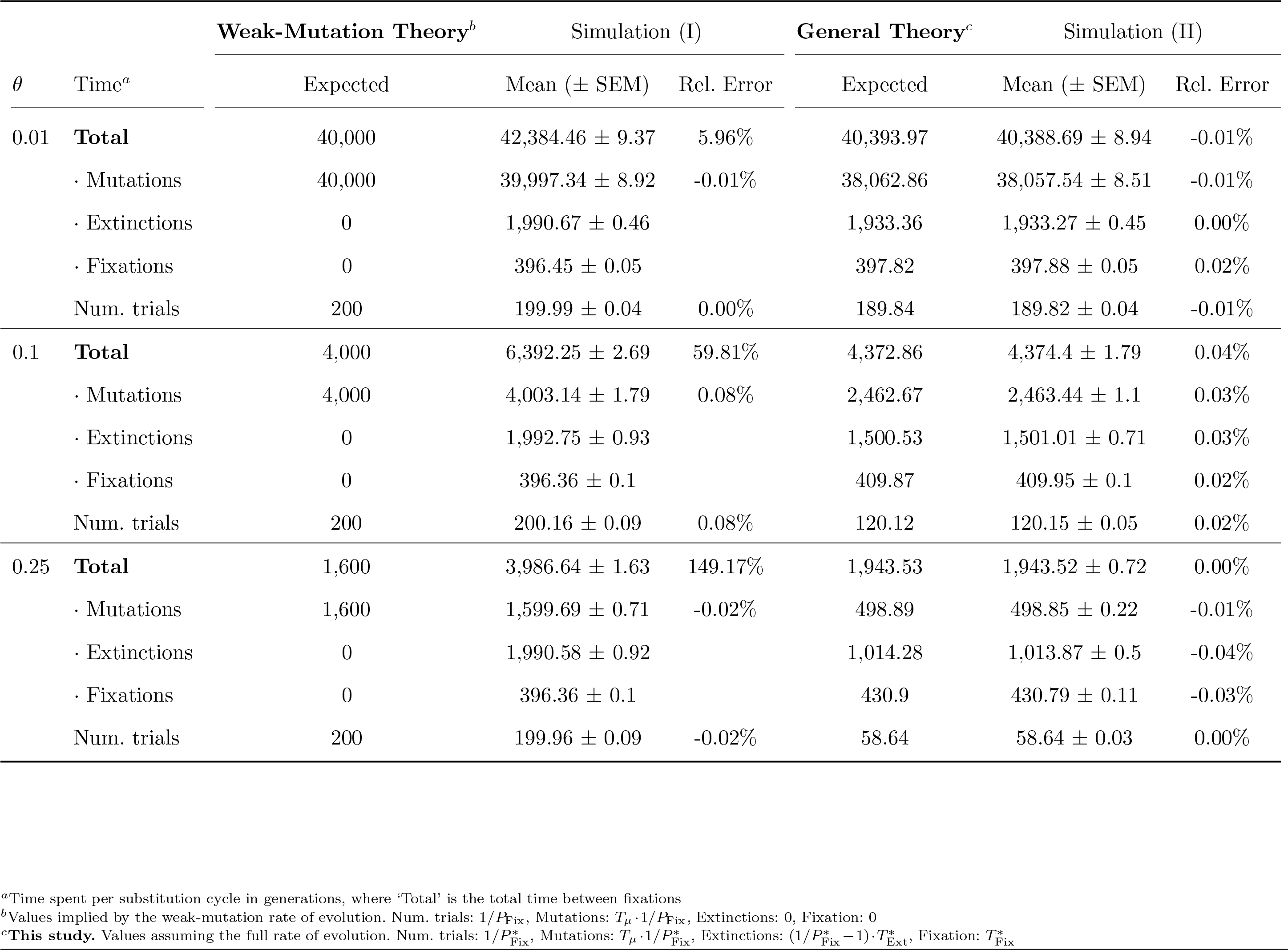
The rate of neutral evolution is slower than the mutation rate when *θ* is modestly large. The mean time between fixations is longer than one over the mutation rate, regardless of whether mutations are considered one-at-a-time in a Wright-Fisher (WF) model with Poisson mutation (‘Simulation I’), or are allowed to arise at any time according to their underlying rates in the WF model (‘Simulation II’). A population size of *N* = 100 was assumed for consistency with later simulations that were more computationally expensive. Note, the only way for the rate of neutral evolution (1/Total) to equal the mutation rate is to assume that the time between fixations is completely determined by the waiting time for mutations, even though it is not. Theoretical predictions from the full rate of evolution were consistent with uniform mutation simulations, regardless of *θ*.

For all values of *θ* considered, the average time between fixations (‘Total’) was significantly longer than the time spent waiting for mutations (‘Mutations’). For small *θ*, the two quantities were relatively similar. But for larger population mutation rates (e.g., *θ* = 0.25), the average time between fixations was *>* 2*X* the time spent waiting for mutations. Thus, even in the one-at-a-time model assumed by the weak-mutation rate of evolution, the absorption times are non-trivial for surprisingly small values of *θ*. In fact, they are proportionally even greater than the absorption times when uniform mutation is fully accounted for (Table 1, Simulation II).

In contrast to the weak-mutation rate of evolution, which is not even consistent with the one-at-a-time model it assumes, when Wright-Fisher simulations were performed by allowing mutation to occur at any time (Table 1, Simulation II), simulation averages were very close to their exact predictions computed from equation 4 (’General Theory’) using WFES (Krukov *et al*., 2016). Notably, under both models, the cumulative time spent waiting for extinctions can be even larger than the time spent waiting for mutations when *θ* is moderately large (e.g., when *θ* = 0.25; Table 1, Simulations I and II). Despite that each mutant-state extinction is generally quite fast, the expected number of mutation-extinction cycles per fixation are generally large and thus can have a substantial influence on the rate of evolution if the expected time it takes for a mutation approaches the expected time for a mutant-state extinction (explored below).

### Is the neutral rate of evolution equal to the mutation rate for other definitions of fixation?

When treating general recurrent mutation in the above analysis, we assumed that substitutions leave the population identical-by-state (IBS), where the population may or may not also be identical-by-descent (IBD; i.e., we have assumed an “IBS/IBD” definition of fixation). This assumption is shared by all standard Wright-Fisher models that are modified to account for mutation (e.g., Ewens, 2004), and is required *ipso facto* when accounting for soft selective sweeps. However, a good deal of population genetic theory instead defines fixation as leaving the population IBD for a single mutant lineage (although as we have argued, this is often done merely by just ignoring new mutations, as in the one-at-a-time mutation model).

It is therefore worth knowing whether the deceleration in the rate of neutral evolution compared to weak-mutation theory critically depends on the definition of fixation. To address this, we performed individual-based simulations of neutral evolution using SLiM (Haller and Messer, 2017), where we could track lineages of mutations under different definitions of fixation (see Methods). In these simulations, new mutations were allowed to arise at any time according to the mutation rate.

Simulation averages for the time between fixations generally indicated that the rate of neutral evolution is slower than the mutation rate for both IBD and IBS/IBD definitions of fixation (Table 2). As expected by our theory, the difference between the actual rate of neutral evolution and the mutation rate increased for increasingly large population mutation rates. Simulation averages for the time between fixations differed from one over the mutation rate by only about 1% for *θ* = 0.01, but differed by as much as 25% when *θ* = 0.25. Importantly, the average time between IBD fixations was generally even longer than for IBS/IBD fixations (except for the case where *θ* = 0.01, where simulation standard errors were relatively large and the mean time was similar under both definitions). This result can be understood as a consequence of IBD populations also being IBS. Because of this, an IBD fixation cannot occur before the population is IBS. However, an IBD fixation can happen after the population becomes IBS. Thus, IBD fixations are expected to take at least as long on average, or longer, than IBS fixations. These results suggest that the noted deceleration in the rate of evolution with increasing mutation rates is not critically dependent on how fixation is defined.

**Table 2.**
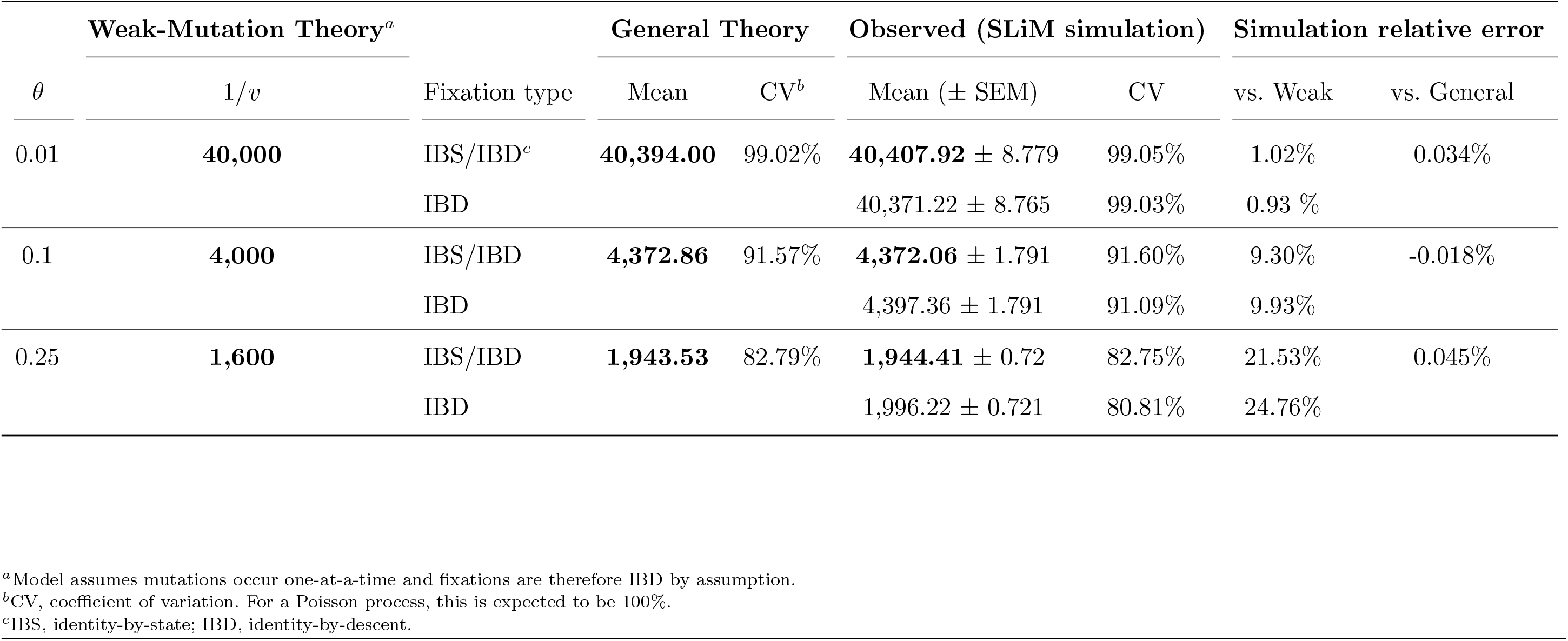
The time between neutral fixations deviates from weak mutation theory for both IBS/IBD and IBD definitions of fixation when mutation is recurrent and *θ* is modestly large. The deceleration effect caused by the segregation times applies to both IBD and IBS/IBD definitions of fixation. For IBS/IBD fixations, our theory and simulations agree precisely, but depart from standard, weak-mutation theory for larger values of *θ*. In all cases, a population size of *N* = 100 was used to make simulations tractable; even for such a small population, simulations took many CPU days to run. Exact predictions were made by directly computing the expected time between fixations and its variance with WFES (Krukov *et al*., 2016), which took less than one second. The number of mutational origins for IBS/IBD fixations are shown in Table S1.

### Implications for the rate of non-neutral evolution

Based on equation 4, when mutation is weak 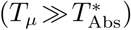, the time between fixations should be dominated by the time spent waiting for mutations (Fig. 1A; as first recognized by Kimura, 1983). However, when mutation is not weak (Fig. 1B), the cumulative time mutants spend segregating in the population before a fixation can become significant, causing the weak mutation model to underestimate the time between fixations and therefore to overestimate the substitution rate (Fig. 1B). This effect can be directly observed for the rate of non-neutral evolution by numerically comparing the weak-mutation substitution rate and the full substitution rate under Wright-Fisher assumptions (Fig. 1C), where the overestimation by the weak mutation model is found to be exaggerated by increasing positive selection. This exaggeration is especially concerning because it implies that methods comparing the rates of substitution for neutral and non-neutral changes (e.g., Goldman and Yang, 1994; McDonald and Kreitman, 1991; Nielsen and Yang, 2003; Zhang *et al*., 2005) are subject to systematic error without correcting for the deceleration effect. These results were replicated and validated in two types of simulations (Fig. 1C; also see Fig. S4, and Methods).

**FIG. 1.**
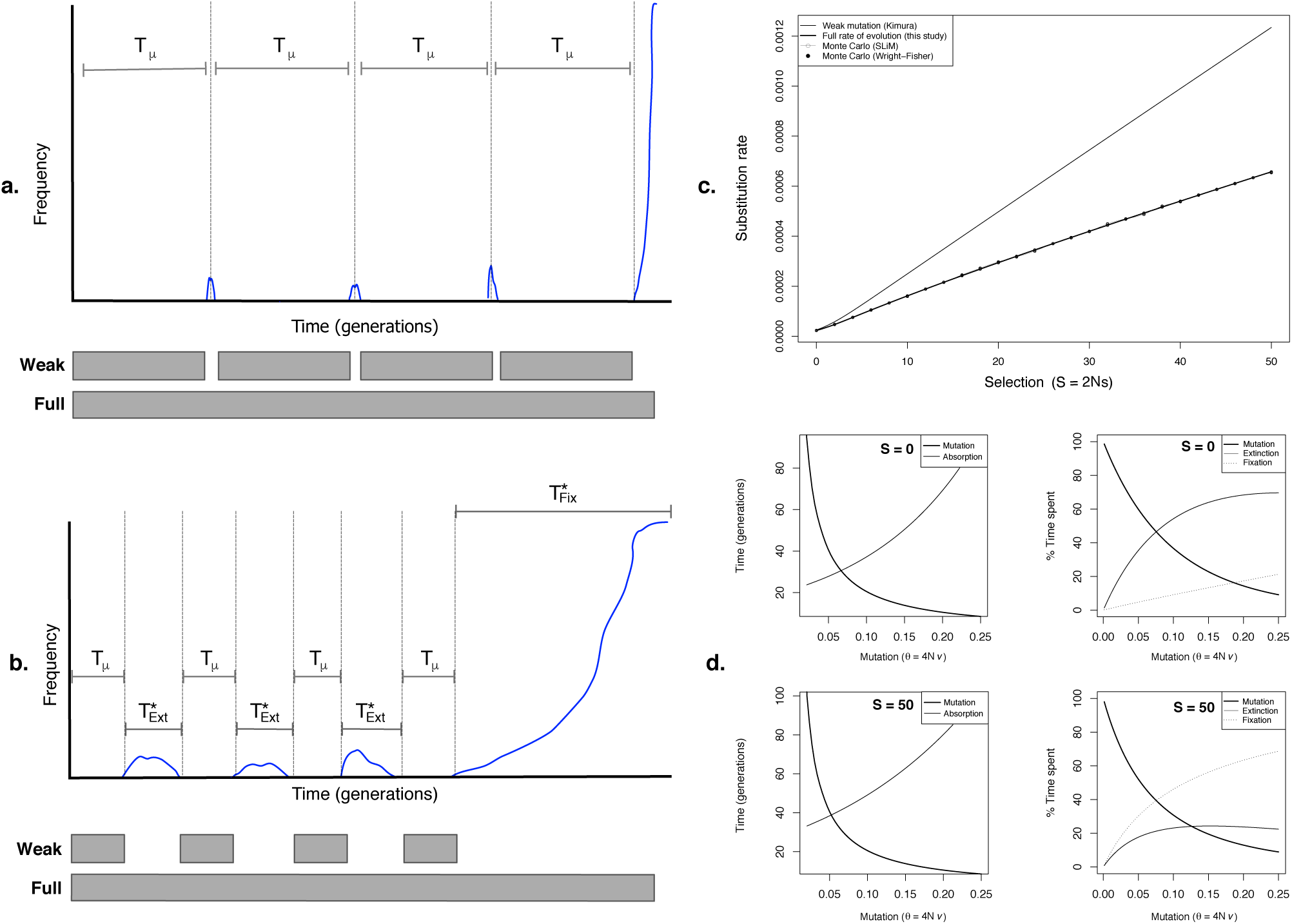
The rate of evolution when mutation may not be weak. **A.** When mutation is weak, the time spent waiting for mutations dominates the time between fixations. **B.** When mutation is not weak, the segregation times also become important. **C.** Demonstration and validation of a deceleration in the rate of evolution under the Wright-Fisher model (*θ* = 0.1, *h* = 0.5, *u* = 0; where *u* is the backward mutation rate). Monte Carlo simulations measured the average number of fixations per generation for a large number of absorptions (SLIM: 5M generations, averaged over 5 runs; Wright-Fisher: 10M absorptions, averaged over 100 runs). Note that both simulations and the full rate of evolution produced nearly identical results, which are fully overlapping. At the origin (*S* = 0), the weak-mutation and full rates of evolution differ by about 10%. **D.** Left: Mean time to mutation (*Tµ*) and absorption 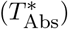. Right: Fraction of the time between fixations spent waiting for mutations, extinctions, and fixations. As explained in the main text, extinction and fixation do not necessarily refer to the fate of a particular lineage of segregating mutants, but to the frequency of the mutant state.

To provide some intuition about the deceleration effect, the mean time spent waiting for a mutation or an absorption is shown in Fig. 1D (left) for increasing population mutation rates. Remarkably, the mean time per absorption for neutral variants overtakes the mean time to a mutation when *θ* is as small as 0.07, strongly violating Kimura’s second weak mutation assumption (see Fig. S5 for a larger parameter range). When *θ* exceeds this value, the rate of neutral evolution is dominated by the time mutants spend segregating in the population prior to a mutation-fixation cycle (Fig. 1D, upper right, “Extinctions”; also see Fig. S6). For selected variants, *T*_*µ*_ is overtaken by 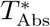 at even smaller values of *θ* (e.g., *θ* = 0.05, Fig. 1D, bottom left). Interestingly, when positive selection is strong, the rate of evolution is dominated by the time it takes for a mutant, once arisen, to go to fixation (Fig. 1D lower right, “fixation”; also see Fig. S6). These observations conspicuously contradict Kimura’s oft-repeated claim that the time it takes for variants to reach their fates does not affect the rate of evolution (Kimura, 1983; Kimura and Ota, 1971). They also clarify that this claim was justified only by the assumption that mutation is always weak.

Examination of the joint effect of mutation, selection, and dominance on the two rates of evolution shows that the deceleration effect in Fig. 1C becomes exaggerated as mutation, selection and dominance are increased (Fig. 2A). Color corresponds to the full rate of evolution as a percentage of the weak-mutation rate of evolution, including dominance where appropriate. As can be more clearly observed in Fig. 2B, for modestly high but biologically realistic values of *θ* (e.g., *θ>* 0.05), a deceleration in the neutral molecular clock (*S* = 0) is predicted when compared to the weak mutation model (as observed in Tables 1 and 2). In fact, the rate of neutral evolution becomes dependent on the population size such that greater decelerations are observed as population mutation rates are increased. This surprising result means that even for strictly neutral mutations, the rate of the molecular clock is expected to be erratic over time if any lineages grow large enough in terms of their population mutation rates. It should also be noticed that nearly neutral variants (*S* = 2) experience a significantly larger deceleration than do neutral variants, which may be consequential for the molecular clock when functional or constrained sequences are used for dating divergence events. These results thus have the potential to help resolve some persistent paradoxes, for instance, where mutation rates from pedigrees and phylogenetic substitution rates unexpectedly differ (Moorjani *et al*., 2016).

**FIG. 2.**
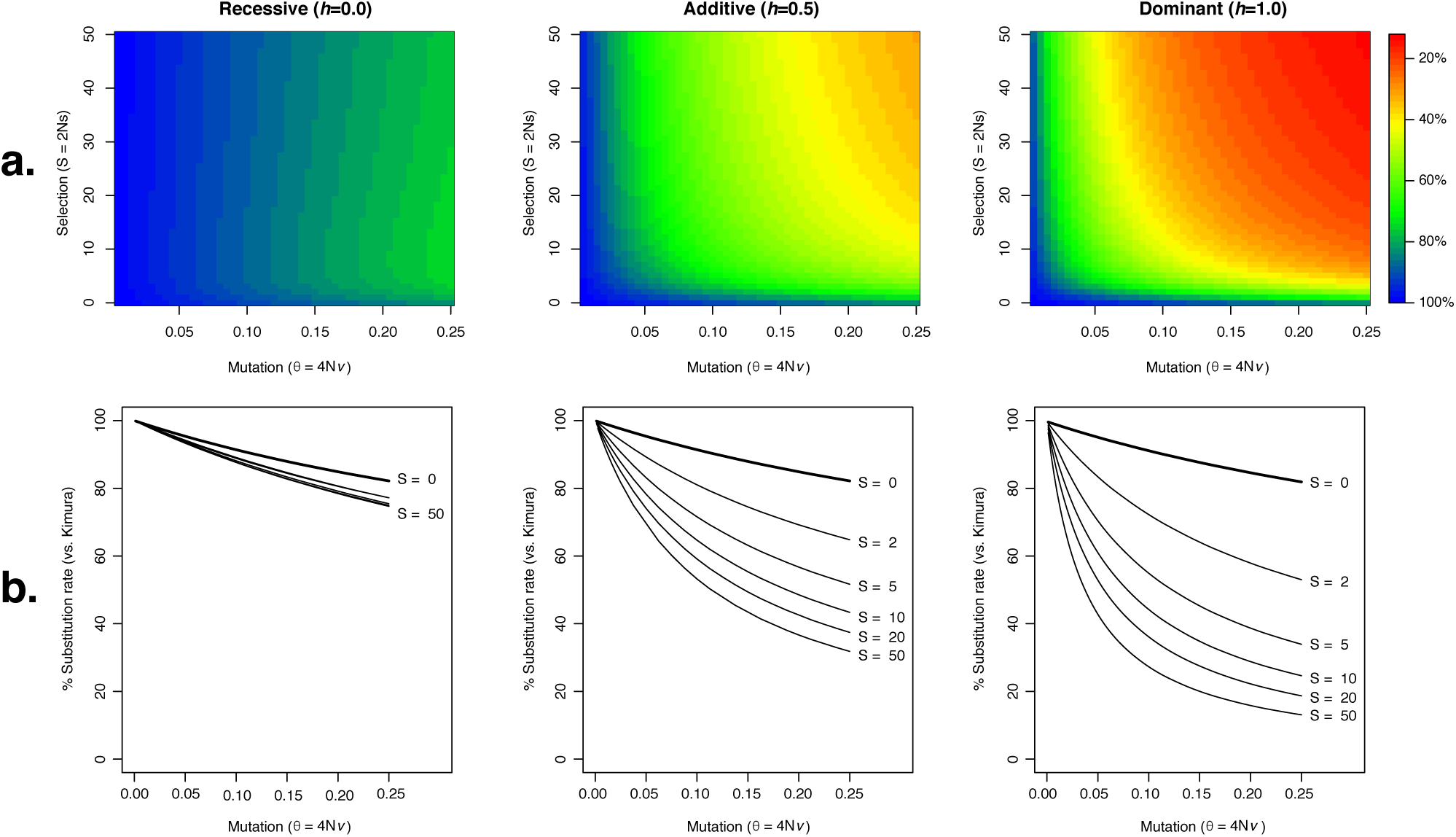
Effect of mutation and selection on the rate of evolution. The full rate of evolution is displayed as a fraction of Kimura’s weak-mutation rate of evolution. **A.** Overall effect of mutation, selection, and dominance across a fine grid. **B.** Specific effect of mutation, selection, and dominance across a range of biologically relevant values.

### The relationship between *d*_*N*_ /*d*_*S*_ (*ω*) and selection unexpectedly depends on the population mutation rate

Since the deceleration effects we observed affect neutral and non-neutral rates of evolution differently, methods that infer the effects of selection in across-species sequence comparisons may be misled if weak-mutation theory is inappropriately applied when *θ ≥* 0.1. One such method, which uses the ratio of non-synonymous to synonymous substitution rates, or *d*_*N*_ */d*_*S*_, is widely used to measure the strength and direction of selection in protein-coding genes. Evidence of heterogeneity in *d*_*N*_ */d*_*S*_ across the lineages of a phylogeny is usually interpreted as evidence of fluctuating or episodically varying selective constraints over time. Contrariwise, we find that when mutation is not weak, adaptive substitutions are expected to have different true *d*_*N*_ */d*_*S*_ values under different population mutation rates, even while the population-scaled selection coefficient is held constant (Fig. 3A). When *θ* is small, *d*_*N*_ */d*_*S*_ increases approximately linearly with population-scaled selection coefficients, with a slope close to 1 for adaptive substitutions (Fig. 3A). However, for larger values of *θ*, the slope of this relationship decreases substantially. Consequently, *d*_*N*_ */d*_*S*_, has different meanings with respect to the actual strength of selection in populations with different population mutation rates. This phenomenon has the potential to make comparison of *d*_*N*_ */d*_*S*_ between species problematic whenever *θ* is large and varies between lineages. It should be noted that while these results may appear superficially similar to earlier results on the population genetics of *d*_*N*_ */d*_*S*_ when unfixed polymorphisms are used to approximate divergence (Kryazhimskiy and Plotkin, 2008), they seem to us to be unrelated.

**FIG. 3.**
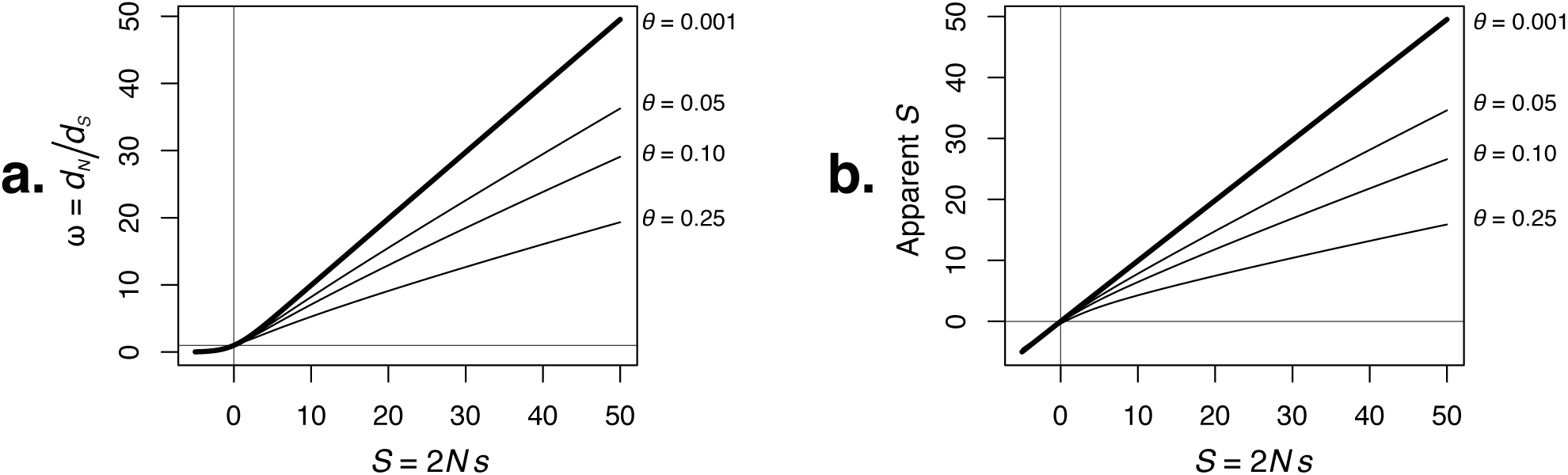
The relationship between population-scaled selection coefficients and measures of selection varies with the population mutation-rate. **A.** The relationship between true *d*_*N*_ */d*_*S*_ and selection is modified by the population mutation rate. The substitution rate was calculated using equation 4 for both *d*_*N*_ and *d*_*S*_ (given the true value of *S*). **B.** *S* inferred under weak mutation assumptions systematically underestimates the true strength of adaptive evolution, particularly for large values of *θ*. Similarly, neutral evolution increasingly appears as weak negative selection for large *θ*.

### Inferred selection coefficients from phylogenetic data become increasingly biased for larger population mutation rates

Across-species population genetics approaches have recently become an important way to make inferences about the relative fitness of different sequence states at the same position (Dimmic *et al*., 2000; Halpern and Bruno, 1998; Nielsen and Yang, 2003; Rodrigue *et al*., 2010; Thorne *et al*., 2007). All such approaches to date use equation 1 and thus assume weak mutation. To determine how the weak mutation approximation might effect inferences about selection when population mutation rates are not small, we numerically solved for the selection coefficient in equation 1 that best approximates the full rate of evolution (using equation 4), across a range of true selection coefficients and population mutation rates. Weak mutation approximations lead to systematic underestimation of the strength of selection, increasingly for larger values of the population mutation rate (Fig. 3B). Importantly, when *S* = 0, inferences of *S*, *Ŝ* (or apparent *S*), suggest increasingly strong negative selection for increasing population mutation rates. For example, for neutral evolution (*S* = 0) and *θ* = 0.1, *Ŝ* under weak mutation assumptions is -0.18. Similarly for *θ* = 0.25, *Ŝ*= -0.38 under weak mutation assumptions. Such findings have the potential to erroneously imply weak constraint in its absence, or perhaps saturation of substitutions. A similar effect is observed for adaptive substitutions, where weak mutation approaches lead to systematically underestimated selection coefficients. For example, at *θ* = 0.1, the strength of strong positive selection (*S* = 50) is inferred using weak mutation calculations as *Ŝ*= 26.6 (53.2% of actual). When theta is larger, *θ* = 0.25, the underestimation is even more extreme, with *Ŝ*= 15.9 (32% of actual). To overcome these problems, the full rate of evolution could simply be used instead of the weak-mutation rate of evolution. More development, however, is required to properly account for mutation to a finite number of alternate alleles in a fully multi-allelic framework (e.g., Wilson *et al*., 2011).

### Limitations of the fixation concept and effect of bidirectional mutation

Equation 4 makes no assumptions about the strength of mutation and should therefore be an accurate expression for one over the mean time between substitutions for all values of *θ*. However, in both single-and multi-locus theory, it is possible that populations with very large *θ* may no longer be able to reach fixation for any one variant in particular (e.g., as implied by Guess and Ewens 1972; Kimura 1983; except perhaps for highly adaptive variants in unchanging selective environments). In this circumstance, the concept of repeated substitutions at the same position could become inadequate to characterize molecular evolution, even if populations continue to visit the fixation boundary at a predictable, but very slow, rate. We therefore sought to determine whether there are any natural limits to the rate of evolution, which can be discerned through single locus theory. For convenience, we will here refer to the population mutation rate parameter for forward mutation as *θ*_*F*_, and the population mutation rate parameter for backward mutation as *θ*_*B*_.

In the previous sections we found that the assumptions of the weak-mutation rate of evolution (equation 1) became strongly violated when *θ ≥* 0.1. To understand how “strong” mutation is in this regime, we computed expected properties of substitution cycles using a modified version of WFES, assuming the Wright-Fisher model and values of *θ* spanning a large range (Table 3). When *θ* = 0.1, more than half of the substitution cycle is expected to be spent with an allele frequency of zero (56.32%), and the average allele frequency throughout the cycle is only 8.53%. In this regime, populations are also expected to spend 80.24% of the time at an allele frequency under 10%. Therefore, we are clearly not in the regime where mutation should be considered “strong” (despite that certain weak-mutation assumptions are violated). Simpler thinking also supports this conclusion, since *θ* between 0.05 and 0.1 implies that a new mutation arises in the population only every 20-40 generations on average. We refer to values of *θ* in this range as “moderate”.

**Table 3.** Statistics of expected substitution trajectories illustrate that faster mutation results in faster neutral evolution when mutation is unidirectional (*θ*_*F*_ = 4*Nv*; *θB* = 0). As before, ‘total time’ refers to the mean time between fixations, which is the reciprocal of the rate of evolution.

| $\theta$ | Total time | $1/v$ | $E[AF]^a$ | Expected fraction of time per substitution cycle when: | | | | |
| --- | --- | --- | --- | --- | --- | --- | --- | --- |
| | | | | $AF=0$ | $AF < 10\%$ | $AF < 25\%$ | $AF < 50\%$ | $AF < 75\%$ |
| 0.0001 | 4,000,396.4 | 4,000,000 | 0.0001 | 0.9994 | 0.9998 | 0.9999 | 0.9999 | 1.0000 |
| 0.001 | 400,396.3 | 400,000 | 0.0010 | 0.9941 | 0.9977 | 0.9986 | 0.9993 | 0.9997 |
| 0.01 | 40,394.0 | 40,000 | 0.0098 | 0.9423 | 0.9772 | 0.9863 | 0.9932 | 0.9972 |
| 0.1 | 4,372.9 | 4,000 | 0.0853 | 0.5632 | 0.8024 | 0.8779 | 0.9380 | 0.9742 |
| 0.25 | 1,943.5 | 1,600 | 0.1768 | 0.2567 | 0.5981 | 0.7408 | 0.8642 | 0.9425 |
| 0.5 | 1,105.9 | 800 | 0.2766 | 0.0828 | 0.3940 | 0.5843 | 0.7719 | 0.9008 |
| 0.75 | 810.8 | 533.3 | 0.3422 | 0.0333 | 0.2784 | 0.4793 | 0.7022 | 0.8672 |
| 1 | 655.0 | 400 | 0.3893 | 0.0168 | 0.2085 | 0.4047 | 0.6466 | 0.8386 |
| 2 | 397.1 | 200 | 0.4963 | 0.0056 | 0.1003 | 0.2504 | 0.5011 | 0.7520 |
| 5 | 205.5 | 80 | 0.6107 | 0.0054 | 0.0526 | 0.1403 | 0.3302 | 0.6039 |
| 10 | 122.9 | 40 | 0.6745 | 0.0082 | 0.0411 | 0.1059 | 0.2499 | 0.4892 |

When considering limits to the rate of evolution under strong mutation, it should be noted that for unidirectional forward mutation, neutral substitutions will always occur given enough time, and larger population mutation rates (*θ*_*F*_) simply mean that substitutions will occur more rapidly (Table 3). This, however, is not true under bidirectional mutation (Table 4). For equal forward and backward mutation rates, increasingly large *θ* reduces the time between both neutral and adaptive substitutions as expected, but at some critical point, *θ*^*\**^, substitutions stop getting faster and begin to slow down (Fig. 4, see inset).

**Table 4.** Statistics of expected substitution trajectories when mutation is bidirectional (*θ* = *θF* = *θB*). Note that the behaviour of the rate of neutral evolution differs much more substantially from 1/*v* than with only forward mutation (c.f., Table 3). ^*a*^Mean expected frequency of the mutant state during the substitution cycle.

| $\theta$ | Total time | $1/v$ | $E[AF]^a$ | Expected fraction of time per substitution cycle when: | | | | |
| --- | --- | --- | --- | --- | --- | --- | --- | --- |
| | | | | $AF=0$ | $AF < 10\%$ | $AF < 25\%$ | $AF < 50\%$ | $AF < 75\%$ |
| 0.0001 | 4,000,792.9 | 4,000,000 | 0.0001 | 0.9994 | 0.9998 | 0.9999 | 0.9999 | 1 |
| 0.001 | 400,793.3 | 400,000 | 0.001 | 0.9941 | 0.9977 | 0.9986 | 0.9993 | 0.9997 |
| 0.01 | 40,795.6 | 40,000 | 0.0098 | 0.9423 | 0.9772 | 0.9863 | 0.9932 | 0.9972 |
| 0.05 | 8,807.1 | 8,000 | 0.0458 | 0.7463 | 0.8932 | 0.935 | 0.9673 | 0.9865 |
| 0.1 | 4,823.8 | 4,000 | 0.0854 | 0.5627 | 0.802 | 0.8777 | 0.9379 | 0.9742 |
| 0.25 | 2,490.9 | 1,600 | 0.1788 | 0.2534 | 0.5931 | 0.7369 | 0.8623 | 0.9423 |
| 0.5 | 1,878.7 | 800 | 0.2871 | 0.0755 | 0.3722 | 0.5642 | 0.7607 | 0.8985 |
| 0.75 | 1,956.0 | 533.3 | 0.3637 | 0.0249 | 0.2367 | 0.4358 | 0.6771 | 0.8634 |
| 1 | 2,463.0 | 400 | 0.4188 | 0.0088 | 0.1506 | 0.3386 | 0.6098 | 0.8387 |
| 2 | 20,347.1 | 200 | 0.4951 | 0.0002 | 0.03 | 0.1597 | 0.504 | 0.8456 |
| 5 | 64,350,775.6 | 80 | 0.5 | 0 | 0.001 | 0.0486 | 0.4939 | 0.9475 |
| 10 | 4,417,890,000,000.0 | 40 | 0.5 | 0 | 0 | 0.0093 | 0.4913 | 0.9893 |

**FIG. 4.**
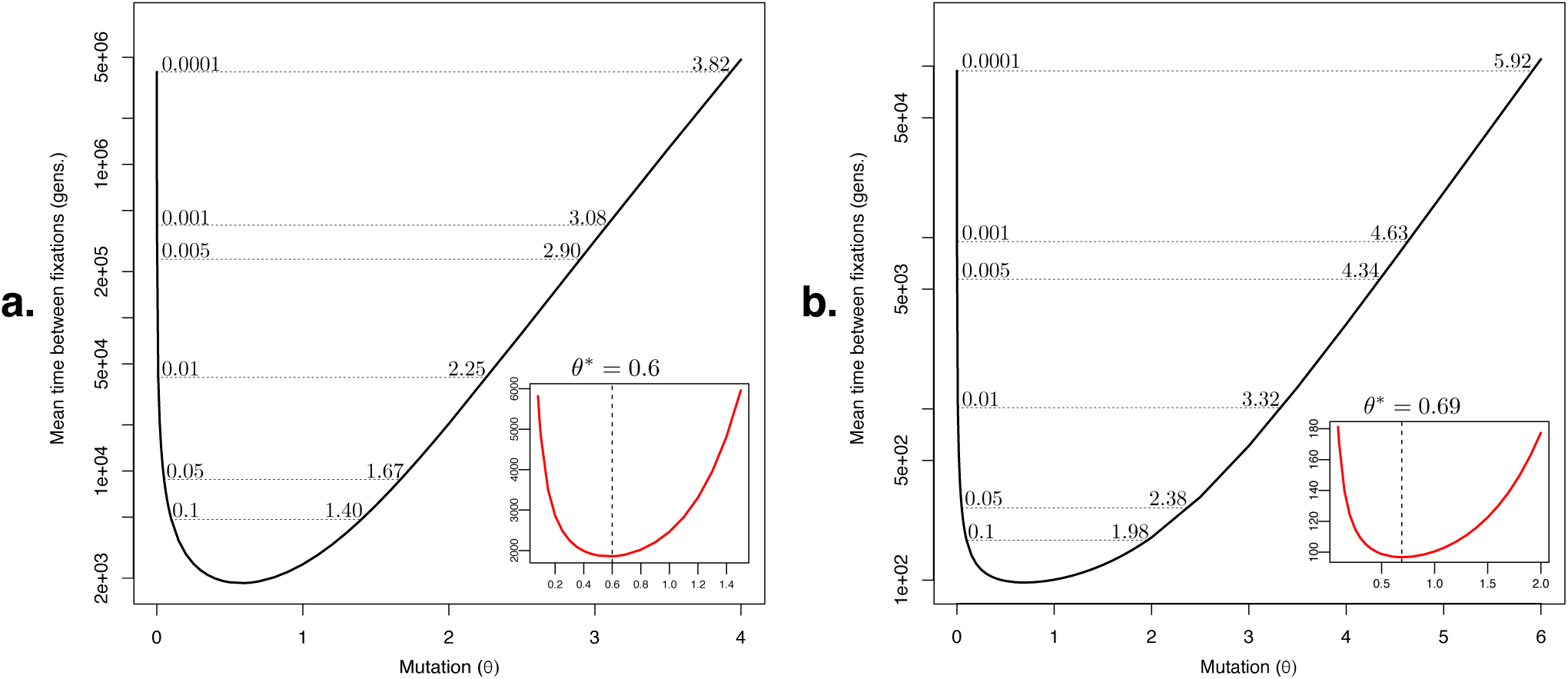
The rate of molecular evolution slows down under strong bidirectional mutation (*θ* = *θ*_*F*_ = *θ*_*B*_), and is maximized at moderate values of *θ*. Isoclines represent equal rates of molecular evolution for small and large *θ* (compare left versus right). **A.** Neutral evolution. We tentatively define: weak mutation (to *θ* = 0.07; see main text), moderate mutation (0.07 *< θ ≤* 0.6), strong mutation (0.6 *< θ ≤* 3.82), and fixation-unlikely (*θ>* 3.82). Inset: point of inflection, *θ*^*\**^ = 0.6, which is the mutation rate that maximizes the rate of neutral evolution under single locus theory. **B.** Strong positive selection (2*Ns* = 50). We tentatively define: weak mutation (to *θ* = 0.05; see main text), moderate mutation (0.05 *< θ ≤* 0.69), strong mutation (0.69 *< θ ≤* 5.92), and fixation-unlikely (*θ>* 5.92). Inset: point of inflection, *θ*^*\**^ = 0.69, which is the mutation rate that maximizes the rate of adaptive evolution.

For very large values of *θ*, fixation requires so much time that it is no longer likely to occur at all (e.g., when *θ*_*F*_ = *θ*_*B*_ = 10, neutral fixations are expected only every 4 trillion generations, Table 4; *N* = 100).

What range of *θ* is then compatible with molecular evolution being driven by successive substitution events? In order to introduce some conservative bounds on how large a rate of population mutation might be considered compatible with evolution by successive substitutions, isoclines can be drawn in Fig. 4 to represent rates of evolution that are equivalently obtained with small and large *θ*. For example, the human population mutation rate parameter has been estimated to be close to *θ* = 0.001 (Kang and Marjoram, 2011; Sung *et al*., 2012), so we can suppose that large values of *θ* that give the same rate of evolution as when *θ* = 0.001 are still in the regime where fixation is both possible and likely (despite that the dynamics of substitution trajectories are expected to be quite different for small versus large *θ*). For neutral variants and bidirectional mutation, this equivalence is obtained at *θ* = 3.08 (Fig. 4A). Therefore, at values of *θ* up to 3.08, the rate of neutral evolution will be no slower than it is in humans with *θ* = 0.001. For strong positive selection, this equivalence leads to an even higher upper bound of *θ* = 4.63 (Fig. 4B). It is therefore reasonable to conclude that molecular evolution proceeds by successive substitutions up to at least *θ* between 3.08-4.63 (under single locus theory). Indeed, if we think *θ* = 0.0001 is a biologically reasonable value, the upper limit to population mutation rates according to this thinking can be as high as *θ* = 3.82-5.92 (Fig. 4). An important caveat to these results is that they depend on single-locus, biallelic theory with equilibrium demography, and the critical value may therefore change when considering multiallelic, multilocus evolution and/or non-equilbrium demographic processes.

An interesting consequence of the point of inflection in Fig. 4B is that under single locus theory, an “optimal” mutation rate exists, which maximizes the rate of adaptive evolution (i.e., *θ* = 0.69 in Fig. 4B, for 2*Ns* = 50, and between 0.6 and 0.69 for weaker positive selection; Fig. 4A). We believe this finding is novel, and it suggests a natural limit to the utility of increasing mutation rates in populations that are required to evolve very rapidly, such as in RNA viruses.

## Discussion

Classical population genetic theory is a source of deep insight into a variety of critical problems in the post-genomic era. However, classical results were built upon assumptions that may now be questioned in the light of recently accumulated knowledge. Many species seem to have population mutation rates substantially below *θ* = 0.1, and in these cases, the use of the classical weak mutation assumption appears unproblematic. Nevertheless, it is not unusual for estimates of *θ* to exceed 0.1, as is true in some hyperdiverse eukaryotes (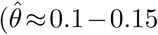 Cutter *et al*. 2013), many prokaryotes (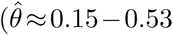 e.g., *Helicobacter pylori*, Sung *et al*. 2012; *Salmonella enterica*, Sung *et al*. 2012; *Pseudomonas syringae*, Hughes *et al*. 2008), pathogens including Plasmodium (a protist, 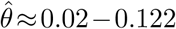 Anderson *et al*. 2017), many DNA viruses (median 0.02, with 22% *≥* 0.1; *n* = 76, censored at 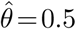, Hughes and Hughes 2007); many to most RNA viruses (median 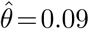, with 49% between 0.1 and 0.5, *n* = 146, censored at 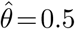, Hughes and Hughes 2007); and HIV-1 (with some estimates of *θ>* 1; Maldarelli *et al*. 2013; Pennings *et al*. 2014; Rouzine *et al*. 2014). Furthermore, since most such estimates are made using the infinite-sites assumption, which forbids recurrent mutation, and since they do not account for background selection on synonymous sites (e.g., when using measures of synonymous diversity to approximate *θ*), they are expected to be conservative and to thus reflect an underestimation of population mutation rates in general. The population mutation rate can be similarly large, or larger, for mutation types with faster natural rates, such as sequence transitions in some organellular genomes, microsatellite mutations, simple repeat polymorphisms, and epigenetic variations. Even in vertebrate mitochondrial DNA, which is not a particularly extreme example, there is evidence of *θ* falling in the range of 0.1-0.3 in many species (Piganeau and Eyre-Walker, 2009). Our findings suggest that in such cases, ignoring the full effect of mutation is unwise.

Kimura and coworkers clearly understood that the equality of the rate of neutral evolution and the mutation rate was subject to some weak-mutation assumptions, which they mentioned in several places (e.g., Crow and Kimura 1970, p. 369; Kimura 1983, p. 46; and Kimura and Ohta 1973). It should therefore not be surprising that the rate of neutral evolution deviates from the mutation rate when mutation is strong, although we feel it is surprising that it deviates so much from weak-mutation theory even for only modestly large values of *θ*. Our results differ from Kimura’s at intermediate population mutation rates mainly because the time it takes variants to reach their fates sets a previously unappreciated fundamental speed-limit on the rate of evolution, which exists, but is of minor consequence when mutation is very weak. As mutation becomes stronger, this speed-limit remains, but becomes more important as the time spent waiting for mutations approaches and then falls below the time it takes for variants to reach their fates. This interpretation also explains why stronger positive selection and dominance lead to greater deviations in the rate of evolution from the weak-mutation theory (Fig. 2).

Despite that it has been claimed by Crow and Kimura (1970, p. 369) that the second weak mutation assumption only requires *θ ≪* 1.0, it does not appear to have been appreciated just how much smaller than 1 the population mutation rate needs to be for mutation to be accurately considered weak. For example, Kimura and Ohta (1973) gave an example of mutant trajectories representing two successive neutral substitution cycles with *θ* = 0.2 (their Figure 1). While this example was labelled as if it conformed exactly to weak mutation assumptions, it is clear even in their figure that the time to fixation, 4*N*_*e*_, was not negligible compared to the time spent waiting for mutations per fixation, 1/*v*. Indeed, our own results show that in this parameter range, the weak-mutation rate of evolution is off by between 60% and 150% compared to simulations under the assumed one-at-a-time mutation model (Table 2).

Our general approach is not without precedent. During the preparation of this manuscript, we became aware of the largely overlooked study by Guess and Ewens (1972), which aimed to determine the effect of mutation on the rate of evolution using an infinite alleles approach. That approach was criticized by Kimura (p. 47 of Kimura 1983), owing to the presumed inappropriateness of modelling substitution processes with infinite alleles. Nevertheless, the spirit of their approach is captured by including mutation in the transition matrix of the underlying model as we have done, and by then approximating the time between fixations as 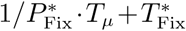, which follows from their equation 22. Note that this approach differs from the full rate of evolution by ignoring the expected time populations spend segregating variants in unsuccessful trials that precede a mutation-fixation cycle. Even though each failed trial generally happens quickly, cumulatively, this time can be very large per expected fixation (e.g., Fig. 1D, right). As shown in Fig. S1, ignoring this fact leads to wild overestimation of the rate of evolution (where the Guess and Ewens model is implemented as described above, in a biallelic context with forward mutation). It is somewhat remarkable that Guess and Ewens also concluded that the weak-mutation rate of evolution overestimates the true rate when *θ>* 0.1. They reached this conclusion, however, for a different reason than we did. In their calculations, they assumed a piecewise biallelic model to approximate infinite alleles. In the biallelic context, however, this model had no forward mutation and included only back mutation (their equation 1). The deceleration in the rate of evolution they inferred is therefore likely a trivial consequence of the slowdown expected by back mutation making it harder for an initial allele to escape extinction. We therefore believe that both studies identified the same critical threshold (*θ* = 0.1) by coincidence.

An advantage of our approach over alternative formulations is that equation 4 is exact and makes no additional assumptions or approximations beyond those of the model used to compute its quantities. Therefore, it remains valid as an expression for one over the expected time between fixations for all values of *θ*. However, as we have pointed out, this is not to say that for some values of *θ*, the meaning of this quantity won’t become incoherent (when mutation is bidirectionally fast; e.g., Table 4). If mutation is strong enough, for example, it may no longer make sense to characterize evolution as being driven by successive fixations at all. This highlights that even the concept of ‘fixation’ itself assumes that mutation is weak to some extent, and that evolution is accordingly mutation-limited. When it is not, both short-term and long-term evolution may be better characterized simply by the changes in allele frequencies that occur over time. These are not new ideas. Some authors have implied that fixation may cease to be a useful concept when *θ* is large (e.g., Guess and Ewens 1972; Kimura 1983), and that instead, measures of ‘establishment’ may be more appropriate (Messer and Petrov, 2013). However, establishment is a mostly subjective concept, and the values of *θ* for which establishment may be more appropriate than fixation remain uncharacterized. It will be interesting in future work to consider the time between fixations, and perhaps the establishment time, while accounting for strong mutation and selection in multi-site models of adaptive evolution, which so far have been mechanistically informative but limited mostly to weak mutation, weak selection scenarios (e.g., Brunet *et al*., 2008; Desai and Fisher, 2007; Rouzine *et al*., 2003, 2008).

Our model, like others, may be criticized for being overly simplistic, as we did not account for the effect of clonal interference among different allelic types, nor account for the effects of linkage. Future work can extend our approach to fully characterize these effects. Nevertheless, there are good reasons to believe that both forces would exaggerate the effects we identified, further slowing the rate of evolution compared to the predictions of classical theory. Therefore our rate of evolution (equation 4) should be considered as an upper bound on the expected rate of evolution when applied to real sequence data. Another obvious future direction will be to extend this work to account for the effects of non-equilibrium demography and selection on the rate of evolution. This may be particularly important in species having average population mutation rates that are low, but that experience periods of large population size (e.g., in Drosophila; Karasov *et al*., 2010).

### Conclusions

1. Our findings suggest that several of the most fundamental rules of molecular evolution fail to generalize when population mutation rates grow beyond a critical threshold that is surprisingly small, regardless of whether mutation is unidirectional or bidirectional. This is true both because, for modestly large population mutation rates: A) the time variants spend segregating in the population before a fixation can become as long, or longer, than the cumulative time spent waiting for mutations (per fixation); and B) the weak-mutation rate of evolution ignores the possibility of soft sweeps, which become more prevalent when *θ* is larger. Through the lens of standard weak-mutation theory, the deceleration effects we identified would be interpreted as either weak negative selection when there is none, as weak positive selection when it is actually strong, or potentially as saturation–a phenomenon where branch lengths can be underestimated due to information loss following many recurrent substitutions that happened in serial over long evolutionary distances. Violations of the predictions of weak-mutation theory are expected when population mutation rates are very large, however, we have identified substantial deviations for modest, biologically relevant population mutation rates (e.g., *θ ≥* 0.1).

2. When recurrent bidirectional mutation is considered, several additional features of the rate of evolution emerge. First, there exists an optimal mutation rate that maximizes the rate of adaptive molecular evolution in Wright-Fisher populations, 0.6 *< θ*^*\**^ *≤* 0.69 (depending on the strength of selection considered). Second, the rate of evolution actually slows down (not decelerates) with increasing bidirectional population mutation rates in excess of *θ*^*\**^. Third, fixation is likely over biologically reasonable timescales for values of *θ* up to between 3 and 6. It will be useful to evaluate the generality of these new results under more realistic models, and to explore whether they can help explain natural variation in mutation rates or population sizes in rapidly-evolving populations.

Taken together, our results suggest that the regime of weak mutation is substantially narrower than has been widely believed. Great caution should thus be exercised in the use of a wide range of evolutionary approaches in organisms with large population mutation rates, such as HIV, hyperdiverse eukaryotes, and many prokaryotes.

## Methods

### Definitions

Throughout this work, and following convention, we refer to the backward mutation rate as *u*, the forward mutation rate as *v*, the dominance coefficient as *h*, and the selection coefficient as *s*. The population mutation rate parameter, *θ*, is defined for diploids as *θ* = 4*Nv*, and the population scaled selection coefficient is defined as *S* = 2*Ns*. We note that the population mutation rate itself is *θ/*2. Since backward mutation will invariably reduce the rate of evolution, and because it is ignored in the weak mutation rate of evolution, we assumed a backward mutation rate of *u* = 0 to be conservative throughout this study, except where otherwise noted.

### Direct computation of the rate of evolution

Let *x* be the current number of mutants in a Wright-Fisher population of size *N*, and *p* the initial number of mutants to arise on a background of *x* = 0. To directly calculate the substitution rate without invoking the theory presented above, we modified our program WFES (Krukov *et al*., 2016) by making *x* = 0 a transient state so that computation of the mean time to absorption from a starting count of zero (*p* = 0) will represent the mean time it takes to go from being 100% wildtype to 100% mutant (i.e., the time between fixations). Similarly, we calculated the variance of the time between fixations as the variance of the time to absorption under these same conditions.

For numerical computations, transition probabilities, *P* (*i*,*j*), were calculated under a Wright-Fisher model including bidirectional mutation, selection, and dominance (Ewens, 2004),

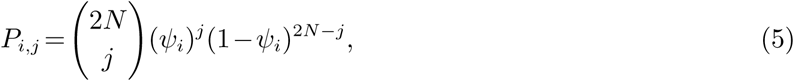

with

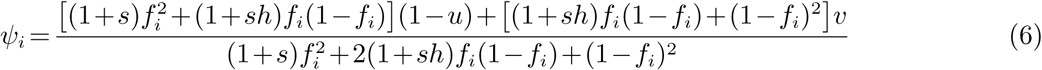

where *f*_*i*_ = *i*/(2*N*). Note that forward and backward mutation are fully accounted for here through their respective rates *v* and *u*.

### Computation of mean times and probabilities

Mean times and probabilities in equation 4 were computed directly from a computationally efficient analysis of the appropriate Wright-Fisher Markov model, using absorbing Markov chain methods we previously described (Krukov *et al*., 2016). To measure properties of expected allele frequency trajectories demarcated by visits to either of the extinction or fixation boundaries, the boundary states were treated as absorbing (except when directly calculating the mean and variance of the time between fixations; see above), even though they may be escaped by mutation. In the general case, this implies that the population evolves until reaching one of the two boundaries, after which it is instantaneously “restarted” by a return process. This return process is equivalent to simply permuting the wildtype and mutant state labels and does therefore not disrupt the behaviour of the model when computing functions of the expected trajectories.

### Simulations

Simulations were performed by three methods so that the effect of variation in the underlying model assumptions or implementations could be examined. First, we simulated directly from the same Wright-Fisher model used to make calculations throughout the manuscript (equations 5 and 6). Populations were assumed to begin as 100% wildtype. For simulations in Table I (Simulation II) and Fig. 1C, The time to the origination of the next founder mutation, or mutations, was drawn from a geometric distribution with success probability, *ρ*, equal to the probability of leaving *x* = 0 under the Wright-Fisher model (with *x* = 0 treated as a transient state).

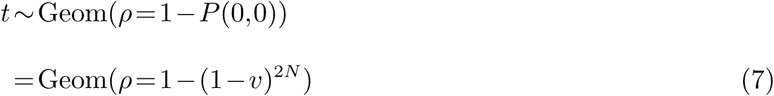

Next, the number of initial mutations, *p*, was drawn from the Wright-Fisher model such that

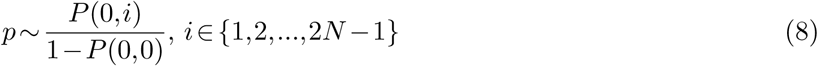

where *P* (0,*i*) is the probability of going from zero copies to *i* copies. The sampling distribution was truncated once the probability of transition became smaller than 10^-8^.

Second, in simulations where mutations were allowed one-at-a-time (Table 1, Simulation I), the same method was used but with some modifications. Mutation rates *u* and *v* were set to zero in equation 6, but were allowed to be non-zero in equations 7 and 8. To make mutations Poisson-distributed, as is assumed by equation 1, *t* was drawn from a Poisson distribution with rate 1/(2*Nv*), and the initial number of mutants *p* was assumed to be 1.

In both of these simulations, the frequencies of the mutant state were then updated iteratively using *P* (*i*,*j*) until either *x* = 0 or *x* = 2*N* were reached. After the population hit either boundary, the simulation was restarted from *x* = 0. Numbers of fixations over many replicates, and the total time spent in generations were recorded. Reported Monte Carlo estimates of the substitution rate were taken to be the number of fixations that occurred divided by the number of generations spent over all simulated absorption cycles. Simulations in Fig. 1 were repeated 100 times and averaged, where each simulation consisted of 10,000,000 absorptions. Simulations in Tables 1 and 2 were run for approximately 5,000,000 fixations, except when *θ* = 0.01, where they were run for 20,000,000 fixations.

Third, for Fig. 1C we performed individual-based simulations using SLiM (Haller and Messer, 2017) under conditions as close as possible to those employed in the first set of simulations. Estimating the Monte Carlo substitution rate as above, estimates of the substitution rate agreed well between the two simulation types. However, for extreme parameter ranges, we found that SLiM slightly over-predicted the substitution rate compared to the Wright-Fisher simulations and the full rate of evolution (Fig. S4). This variation is likely due to minor differences in assumptions or implementation details. Simulation error bars were all very small and appeared as points in Fig. 1C. They were thus omitted from the display item. The Erdos code used to conduct SLiM simulations is included in SI Methods 1.4. Simulations were repeated 5 times and consisted of 5,000,000 generations, where the simulation was restarted after each absorption.

SLiM simulations for Table 2 were performed similarly, but where fixation was varied between IBS/IBD and IBD definitions. For the first definition, the simulations were stopped and restarted when the population reached a frequency of 1.0 for the mutant state, regardless of the number of mutant lineages. For the second, simulations were restarted only when the population reached a frequency of 1.0 for the mutant state from a single mutational lineage. The Erdos code used to conduct these simulations is included in SI Methods 1.5 and 1.6.

## Acknowledgements

The authors gratefully acknowledge Ivan Krukov, Sam Yeaman, Caro-Beth Stewart, David Liberles, Quan Long, and Edwin Wang for comments on the manuscript, and Ben Haller and Phil Messer for help with SLiM. We also thank two anonymous reviewers and Jeff Thorne, for constructive comments that helped us improve the manuscript. This work was supported by a Discovery Grant to APJdK from the Natural Sciences and Engineering Research Council of Canada (NSERC) and by an NSERC undergraduate research award (BDS). The authors gratefully acknowledge infrastructure support from the Canada Foundation for Innovation, Alberta Enterprise and Advanced Education, and the Alberta Children’s Hospital Research Institute.

## Author contributions

APJdK designed the research, developed and implemented the model, performed the analyses, made the figures, and wrote the manuscript. BDS contributed ideas, developed the proof, performed simulations, and edited the manuscript.

The authors declare no conflict of interest.

## References

Allen, B., Sample, C., Dementieva, Y., Medeiros, R. C., Paoletti, C., and Nowak, M. A. 2015. The molecular clock of neutral evolution can be accelerated or slowed by asymmetric spatial structure. PLoS Comput Biol, 11(2): e1004108.

Anderson, T. J., Nair, S., McDew-White, M., Cheeseman, I. H., Nkhoma, S., Bilgic, F., McGready, R., Ashley, E., Pyae Phyo, A., White, N. J., and Nosten, F. 2017. Population parameters underlying an ongoing soft sweep in southeast asian malaria parasites. Mol Biol Evol, 34(1): 131–144.

Brown, W. M., George, M. J., and Wilson, A. C. 1979. Rapid evolution of animal mitochondrial DNA. Proc Natl Acad Sci U S A, 76(4): 1967–71.

Brunet, E., Rouzine, I. M., and Wilke, C. O. 2008. The stochastic edge in adaptive evolution. Genetics, 179(1): 603–20.

Bustamante, C. 2005. Population genetics of molecular evolution. In R. Nielsen, editor, Statistical Methods in Molecular Evolution, pages 63–99. Springer, New York.

Charlesworth, B. and Jain, K. 2014. Purifying selection, drift, and reversible mutation with arbitrarily high mutation rates. Genetics, 198(4): 1587–602.

Crow, J. and Kimura, M. 1970. Introduction to population genetics theory. Harper and Row Publishers.

Cutter, A. D., Jovelin, R., and Dey, A. 2013. Molecular hyperdiversity and evolution in very large populations. Mol Ecol, 22(8): 2074–95.

de Koning, APJ., Gu, W., and Pollock, D. D. 2009. Rapid likelihood analysis on large phylogenies using partial sampling of substitution histories. Mol Biol Evol, 27(2): 249–65.

De Sanctis, B., Krukov, I., and de Koning, APJ 2017. Allele age under non-classical assumptions is clarified by an exact computational markov chain approach. Sci Rep, 7(1): 11869.

Desai, M. M. and Fisher, D. S. 2007. Beneficial mutation selection balance and the effect of linkage on positive selection. Genetics, 176(3): 1759–98.

Dimmic, M. W., Mindell, D. P., and Goldstein, R. A. 2000. Modeling evolution at the protein level using an adjustable amino acid fitness model. Pac Symp Biocomput, pages 18–29.

Draghi, J. A., Parsons, T. L., and Plotkin, J. B. 2011. Epistasis increases the rate of conditionally neutral substitution in an adapting population. Genetics, 187(4): 1139–52.

Ehrlich, M. and Wang, R. Y. 1981. 5-methylcytosine in eukaryotic DNA. Science, 212(4501): 1350–7.

Ewens, W. J. 2004. Mathematical Population Genetics 1: Theoretical Introduction Edn. 2. Springer-Verlag, New York.

Felsenstein, J. 2003. Inferring Phylogenies. Sinauer.

Goldman, N. and Yang, Z. 1994. A codon-based model of nucleotide substitution for protein-coding DNA sequences. Mol Biol Evol, 11(5): 725–36.

Guess, H. A. and Ewens, W. J. 1972. Theoretical and simulation results relating to the neutral allele theory. Theor Popul Biol, 3(4): 434–47.

Haller, B. C. and Messer, P. W. 2017. Slim 2: Flexible, interactive forward genetic simulations. Mol Biol Evol, 34(1): 230–240.

Halpern, A. L. and Bruno, W. J. 1998. Evolutionary distances for protein-coding sequences: modeling site-specific residue frequencies. Mol Biol Evol, 15(7): 910–7.

Hughes, A. L. and Hughes, M. A. 2007. More effective purifying selection on rna viruses than in dna viruses. Gene, 404(1-2): 117–125.

Hughes, A. L., Friedman, R., Rivailler, P., and French, J. O. 2008. Synonymous and nonsynonymous polymorphisms versus divergences in bacterial genomes. Mol Biol Evol, 25(10): 2199–209.

Kang, C. and Marjoram, P. 2011. Inference of population mutation rate and detection of segregating sites from next-generation sequence data. Genetics, 189(2): 595–605.

Karasov, T., Messer, P. W., and Petrov, D. A. 2010. Evidence that adaptation in drosophila is not limited by mutation at single sites. PLoS Genet, 6(6): e1000924.

Kimura, M. 1968. Evolutionary rate at the molecular level. Nature, 217(5129): 624–6.

Kimura, M. 1983. The Neutral Theory of Molecular Evolution. Cambridge University Press.

Kimura, M. and Ohta, T. 1973. Mutation and evolution at the molecular level. Genetics, 73: Suppl 73:19–35.

Kimura, M. and Ota, T. 1971. On the rate of molecular evolution. J Mol Evol, 1(1): 1–17.

King, J. L. and Jukes, T. H. 1969. Non-darwinian evolution. Science, 164(3881): 788–98.

Krukov, I., de Sanctis, B., and de Koning, APJ 2016. Wright-Fisher exact solver (WFES): scalable analysis of population genetic models without simulation or diffusion theory. Bioinformatics, 33(9): 1416–1417.

Kryazhimskiy, S. and Plotkin, J. B. 2008. The population genetics of dn/ds. PLoS Genet, 4(12): e1000304.

Lanfear, R., Kokko, H., and Eyre-Walker, A. 2014. Population size and the rate of evolution. Trends Ecol Evol, 29(1): 33–41.

Lynch, M. 2010. Evolution of the mutation rate. Trends Genet, 26(8): 345–52.

Maldarelli, F. et al. 2013. HIV populations are large and accumulate high genetic diversity in a nonlinear fashion. J Virol, 87(18): 10313–23.

McCandlish, D. M. and Stoltzfus, A. 2014. Modeling evolution using the probability of fixation: history and implications. Q Rev Biol, 89(3): 225–52.

McDonald, J. H. and Kreitman, M. 1991. Adaptive protein evolution at the adh locus in drosophila. Nature, 351(6328): 652–4.

Messer, P. W. and Petrov, D. A. 2013. Population genomics of rapid adaptation by soft selective sweeps. Trends Ecol Evol, 28(11): 659–69.

Messer, P. W., Ellner, S. P., and Hairston, N. G., J. 2016. Can population genetics adapt to rapid evolution? Trends Genet, 32(7): 408–418.

Messier, W. and Stewart, C. B. 1997. Episodic adaptive evolution of primate lysozymes. Nature, 385(6612): 151–4.

Moorjani, P., Gao, Z., and Przeworski, M. 2016. Human germline mutation and the erratic evolutionary clock. PLoS Biol, 14(10): e2000744.

Muse, S. V. and Gaut, B. S. 1994. A likelihood approach for comparing synonymous and nonsynonymous nucleotide substitution rates, with application to the chloroplast genome. Mol Biol Evol, 11(5): 715–24.

Nielsen, R. and Yang, Z. 2003. Estimating the distribution of selection coefficients from phylogenetic data with applications to mitochondrial and viral DNA. Mol Biol Evol, 20(8): 1231–9.

Pennings, P. S. and Hermisson, J. 2005. Soft sweeps: molecular population genetics of adaptation from standing genetic variation. Genetics, 169(4): 2335–2352.

Pennings, P. S. and Hermisson, J. 2006. Soft sweeps II–molecular population genetics of adaptation from recurrent mutation or migration. Mol Biol Evol, 23(5): 1076–84.

Pennings, P. S., Kryazhimskiy, S., and Wakeley, J. 2014. Loss and recovery of genetic diversity in adapting populations of HIV. PLoS Genet, 10(1): e1004000.

Piganeau, G. and Eyre-Walker, A. 2009. Evidence for variation in the effective population size of animal mitochondrial DNA. PLoS One, 4(2): e4396.

Richard, G. F., Kerrest, A., and Dujon, B. 2008. Comparative genomics and molecular dynamics of DNA repeats in eukaryotes. Microbiol Mol Biol Rev, 72(4): 686–727.

Rodrigue, N., Philippe, H., and Lartillot, N. 2010. Mutation-selection models of coding sequence evolution with site-heterogeneous amino acid fitness profiles. Proc Natl Acad Sci U S A, 107(10): 4629–34.

Rouzine, I. M., Wakeley, J., and Coffin, J. M. 2003. The solitary wave of asexual evolution. Proc Natl Acad Sci U S A, 100(2): 587–92.

Rouzine, I. M., Brunet, E., and Wilke, C. O. 2008. The traveling-wave approach to asexual evolution: Muller’s ratchet and speed of adaptation. Theor Popul Biol, 73(1): 24–46.

Rouzine, I. M., Coffin, J. M., and Weinberger, L. S. 2014. Fifteen years later: hard and soft selection sweeps confirm a large population number for HIV in vivo. PLoS Genet, 10(2): e1004179.

Sung, W., Ackerman, M. S., Miller, S. F., Doak, T. G., and Lynch, M. 2012. Drift-barrier hypothesis and mutation-rate evolution. Proc Natl Acad Sci U S A, 109(45): 18488–92.

Thorne, J. L., Choi, S. C., Yu, J., Higgs, P. G., and Kishino, H. 2007. Population genetics without intraspecific data. Mol Biol Evol, 24(8): 1667–77.

Wilson, D. J., Hernandez, R. D., Andolfatto, P., and Przeworski, M. 2011. A population genetics-phylogenetics approach to inferring natural selection in coding sequences. PLoS Genet, 7(12): e1002395.

Wright, S. 1938. The distribution of gene frequencies under irreversible mutation. Proc Natl Acad Sci U S A, 24(7): 253–9.

Yang, Z. 1998. Likelihood ratio tests for detecting positive selection and application to primate lysozyme evolution. Mol Biol Evol, 15(5): 568–73.

Yang, Z. 2014. Molecular evolution: a statistical approach. Oxford University Press.

Yang, Z., Nielsen, R., Goldman, N., and Pedersen, A. M. 2000. Codon-substitution models for heterogeneous selection pressure at amino acid sites. Genetics, 155(1): 431–49.

Zhang, J., Nielsen, R., and Yang, Z. 2005. Evaluation of an improved branch-site likelihood method for detecting positive selection at the molecular level. Mol Biol Evol, 22(12): 2472–9.

